# Neural representations of economic decision variables in human posterior parietal cortex

**DOI:** 10.1101/2023.05.18.541297

**Authors:** Christian Klaes, Artur Pilacinski, Spencer Kellis, Tyson Aflalo, Charles Liu, Richard Andersen

**Affiliations:** Division of Biology and Biological Engineering, California Institute of Technology, Pasadena, CA 91125, USA; USC Neurorestoration Center and the Departments of Neurosurgery and Neurology, University of Southern California, Los Angeles, USA; Ruhr University Bochum, Universitätsstr. 150, 44801 Bochum, Germany; T & C Chen Brain-machine Interface Center at Caltech, USA; Keck School of Medicine, University of Southern California, Los Angeles, USA; CINEICC, University of Coimbra, Portugal

## Abstract

Decision making has been intensively studied in the posterior parietal cortex in non-human primates on a single neuron level. In humans decision making has mainly been studied with psychophysical tools or with fMRI. Here, we investigated how single neurons from human posterior parietal cortex represent numeric values informing future decisions during a complex two-player game. The tetraplegic study participant was implanted with a Utah electrode array in the anterior intraparietal area (AIP). We played a simplified variant of Black Jack with the participant while neuronal data was recorded. During the game two players are presented with numbers which are added up. Each time a number is presented the player has to decide to proceed or to stop. Once the first player stops or the score reaches a limit the turn passes on to the second player who tries to beat the score of the first player. Whoever is closer to the limit (without overshooting) wins the game. We found that many AIP neurons selectively responded to the face value of the presented number. Other neurons tracked the cumulative score or were selectively active for the upcoming decision of the study participant. Interestingly, some cells also kept track of the opponent’s score. Our findings show that parietal regions engaged in hand action control also represent numbers and their complex transformations. This is also the first demonstration of complex economic decisions being possible to track in single neuron activity in human AIP. Our findings show how tight are the links between parietal neural circuits underlying hand control, numerical cognition and complex decision-making.

## Introduction

The process of decision making has been studied intensively in non-human primate (NHP) electrophysiology (Newsome et al., 1989; Platt and Glimcher, 1999; Sugrue et al., 2004; Gold and Shadlen, 2007; Andersen and Cui, 2009; Kable and Glimcher, 2009) and human fMRI studies (Bush et al., 2002; Sanfey et al., 2003; Glimcher and Rustichini, 2004; Hsu et al., 2005; Krain et al., 2006; Martino et al., 2006) where it is also prominently known as neuroeconomics. Several cortical areas seem to be involved in the decision making process and the posterior parietal cortex (PPC) seems to play a critical role (Platt and Glimcher, 1999; Gold and Shadlen, 2007; Rangel et al., 2008; Cisek and Kalaska, 2010; Freedman and Assad, 2011). In former NHP studies it has already been shown that decision variables, like the outcome probability and value, can be extracted from PPC neurons (Platt and Glimcher, 1999; Shadlen and Newsome, 2001; Sugrue et al., 2004). Especially for the lateral intraparietal area (LIP), correlates for perceptual decision making have been found (Shadlen and Newsome, 2001). Neurons in PPC also seem to dynamically adapt to task relevant features and can show ‘mixed selectivity’, which means a combination of many task features to a high level representation (Fusi et al., 2016). However, the complexity of decision making tasks which NHPs can learn with sufficient accuracy is limited by the training time and cognitive abilities of NHPs. Either complex relationships have to be simplified in NHP tasks or a long learning period has to be taken into account. Thus, it is also difficult to compare neuronal behavior in multiple different tasks. Human data from fMRI and EEG studies can partially address these issues but have to deal with low temporal or low spatial signal resolution instead. This, in turn, does not allow capturing the role of single neurons in human decision making, leading to potential difficulties in defining functional areas between humans and NHP such as observed for action planning (for reviews see e.g. Orban, 2016; Foster et al., 2022). Fortunately, in recent years it has become more feasible to obtain electrophysiological data directly from humans since several groups have started pilot studies using brain-machine interface paradigms using electrocorticography (Leuthardt et al., 2004; Shenoy et al., 2008) or implanted electrode arrays (Hochberg et al., 2012; Collinger et al., 2013; Aflalo et al., 2015). Most of the electrode array studies, except Aflalo et al., have targeted the primary motor cortex, focusing on motor action representations. In our own brain-machine interface studies (Aflalo et al., 2015; Klaes et al., 2015) we were able to record data from the parietal cortex while the study participant was engaged in various tasks. In this study, activity of single neurons was recorded through a Utah electrode array implanted in the anterior intraparietal area (AIP). During the recording sessions in which the study participant was primarily learning to control reach and grasp of a robotic limb via brain control we could also integrate various other tasks, thus studying cognitive engagement of neurons tuned to sensorimotor actions. This unique opportunity allowed us to directly track the function of AIP neurons in abstract cognitive tasks such as processing numerosities for upcoming economic decisions. Besides trying to determine whether decision variables are represented in single neuron responses a similar way as has been shown for NHPs (Platt and Glimcher, 1999; Shadlen and Newsome, 2001; Andersen and Cui, 2009), we also wanted to know if neurons in the same region in which we previously recorded neurons selective for sensorimotor aspects of movement trajectories and hand shapes selective neurons (Aflalo et al., 2015; Klaes et al., 2015) would also be active in actions where no motor imagery was instructed but abstract numerical operations had to be performed. As human intraparietal areas have been demonstrated to process numerosities and mathematical operations (Piazza et al., 2004; Knops et al., 2009; Roitman et al., 2012; Kutter et al., 2018), we expected that both numbers and their transformations can be potentially represented in human single neurons activity of AIP, in accordance with prior NHP and human studies (see e.g. Nieder and Miller, 2004, Viswanathan and Nieder, 2013; Kutter et al., 2018).

For the current study we recorded neuronal data while the participant played a two-player press-your-luck game which we used to assess various variables relevant for making decisions in the game. The game featured memory and economic decision making components. Since two players simultaneously participate in the game, we could also observe cognitive processes during the opponent’s turn. We hypothesized that human AIP neurons will represent numeric values for the upcoming economic decisions. Moreover, we expected these representations to be not limited to face values, but also denote complex transformations needed for decision making processes, such as keeping track of the cumulative value. We also expected that neurons would encode the player’s upcoming abstract economic decisions. Lastly, we expected that single neurons in AIP will not only process the subject’s own variables and decisions, but will keep track of those of the opponent.

## Materials and Methods

### Approvals

This study was approved by the institutional review boards at the California Institute of Technology, Rancho Los Amigos and the University of Southern California Los Angeles. We obtained informed consent from the patient before participation in the study. EGS also gave his written consent to be shown in study related videos. We also obtained an investigative device exemption from the FDA (IDE #G120096) to use the implanted devices throughout the study period. This study is registered with ClinicalTrials.gov (NCT01849822).

### Implantation

The subject in this study, EGS, is a male tetraplegic patient who was 32 years old at the time of implantation. His spinal cord lesion was complete at cervical level C3-4 with a paralysis of all limbs. At the time of implantation he was 10 years post lesion. We implanted two 96 channel micro-electrode arrays (Blackrock Microsystems) in two areas of the posterior parietal cortex. One was implanted on the surface of the superior parietal lobule (putative human Brodmann’s Area 5) and one at the junction of the intraparietal sulcus with the postcentral sulcus (putative human anterior intraparietal area, AIP). The exact placement of the arrays was based on an fMRI task which EGS performed prior to implantation. Since the number of units recorded on the AIP array was substantially higher than on the BA5 array we report only on the AIP results. Data sets from the same patient have been used in two other experiments which have already been published (Aflalo et al., 2015; Klaes et al., 2015). The recordings for this study are completely separate from those experiments. Data was collected over a period of 8 months during which other experiments also took place. Details of the implantation are identical to those published by Aflalo (Aflalo et al., 2015) and Klaes (Klaes et al., 2015) and have been repeated here for completeness.

### Placement of electrode arrays and fMRI task

To determine the placement for the two implanted arrays we had the patient perform an imagined hand reaching and grasping task during an fMRI scan.

A GE 3T scanner at the USC Keck medical center was used for scanning. Parameters for the functional scan were: T2*-weighted single-shot echo-planar acquisition sequence (TR = 2000 ms; slice thickness = 3 mm; in-plane resolution = 3 x 3 mm; TE = 30 ms; flip angle = 80; fov= 192 x 192 mm; matrix size = 64 x 64; 33 slices (no-gap) oriented 20 degrees relative to ACPC line). Parameters for the anatomical scan were: GE T1 Bravo sequence (TR = 1590 ms; TE = 2.7 ms; fov =176 x 256 x 256 mm; 1 mm isotropic voxels). Surface reconstruction of the cortex was done using ’Freesurfer’ (http://surfer.nmr.mgh.harvard.edu/).

The reach and grasp task started with a 3 s fixation period in which EGS had to fixate a dot in the center of the screen. In the following cue phase he was cued to the type of imagined action to perform which could be either a precision grip, power grip, or a reach without hand shaping. Next a cylindrical object was presented in the stimulus phase. If the object was ’whole’ (’go’ condition) EGS had to imagine performing the previously cued action on the object and report back the color of the part of the object which was closest to his thumb. The object could be presented in one of six possible orientations and EGS could freely choose how to align his imagined hand with regard to the object. The reported color allowed us to determine if the imagined action was performed with an overhand or underhand posture. Analysis of the orientation of the object and the reported color suggested that EGS was imagining biomechanically plausible, naturalistic arm movements. If the object presented in the stimulus phase was ’broken’ (’no go’ condition) EGS had to withhold the cued action. This was used as a control condition.

The BOLD response from this task was then used to determine possible implantation sites for surgery. Based on the highest activation for grasping and reaching and intraoperative constraints (blood vessel locations, cable and pedestal placement etc.) two implantation sites were picked. Statistical analysis was restricted to the superior parietal lobule to increase statistical power. More details about the fMRI task and its implementation can be found elsewhere (Aflalo et al., 2015). Details of the fMRI pre-surgery task are identical to those published by Aflalo and (Aflalo et al., 2015) and Klaes (Klaes et al., 2015) and have been repeated here for completeness.

### Simple Jack task

Simple jack is a two-player press-your-luck game (Figure 1a). It consists of several game rounds in which each player takes a turn. The game ends when one of the players wins 16 rounds. At the beginning of a turn a text appears on the screen stating the name of the player whose turn it is. This player has to make decisions in this turn and is referred to as the ’active player’. During the active player’s turn a number between 1 and 10 is briefly (500 ms) displayed on the screen. After a short pause (1000 ms) a question mark is displayed. This indicates that the active player has to make a decision. The active player can either choose to ’hit’ or to ’stay’. If the active player chooses ’hit’ another number in the same range will flash on the screen after a short delay (2000 ms). If the sum of all previous numbers and the current number is smaller than 20 (’target number’) the active player will be presented with another decision in which he/she can choose to ’hit’ or ’stay’. If the number is equal or larger than the target number (overshooting) the active player’s turn ends and the next player becomes the active player. If the active player chooses ’stay’ his/her turn ends and the next player becomes the active player. After ending the turn, either through overshooting or by choosing ’stay’, the score for this round is shown for a short delay. After both players take their turn the round ends and the winner for that round is determined by comparing the players’ scores. The player who had the highest sum in his/her turn but did not reach a score higher than the target number wins the round and gains 2 points. In case both players have the same score or both have a score higher than the target number the round results in a draw. In a draw the first player of that round gains 1 point. If one player has a score higher than the target number and the other player is below the target number but higher or equal than 17 (’punishment number’) the player who overshot loses 1 point. The starting player alternated between turns so that EGS would either be the first player to do his turn or the second player. The very first turn of each session would be started by EGS.

**Figure 1:**
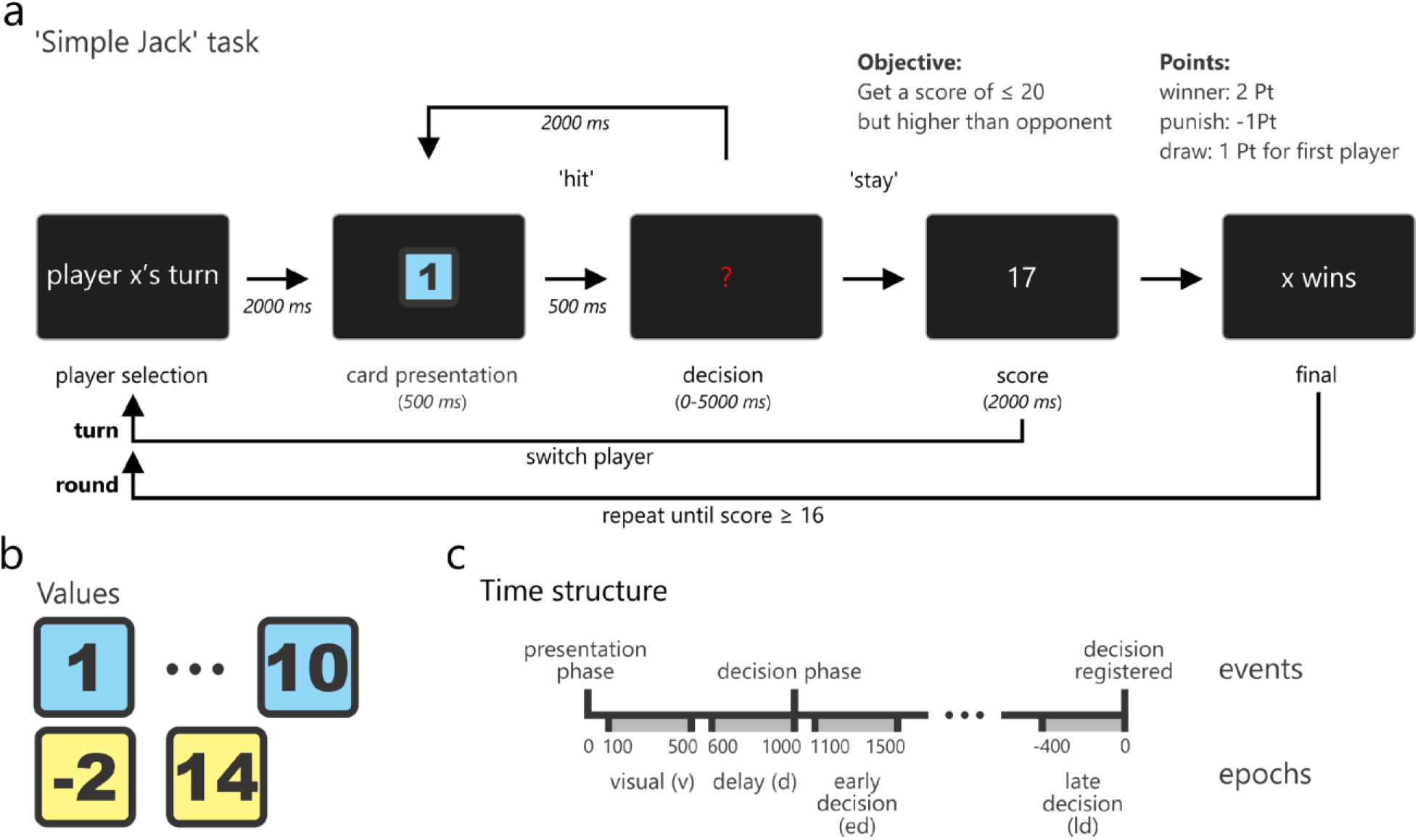
(a) Schematic representation of the general task flow indicating the different phases of the task, their names (below the screen images) and their length (italic numbers). The schematic screen examples show one possible example of events in a task where x would denote the current player who wins in this example with a score of 17. Arrows show the flow of events. The two alternative progressions of the task are indicated by the two arrows labeled ’hit’ and ’stay’. The two long arrows at the bottom indicate all the phases that belong to a turn and a round of the game. (b) Some examples of the card images which were shown to the players. (c) Time structure of the task which also shows the epochs which are used in the statistical analyses. Note that the ’decision registered’ event was the point in time when the experimenter entered the decision of the players which could happen at any time in a 5 s time interval. Times are given in ms and are relative to the presentation phase except for the ’late decision’ epoch, which was defined relative to the ’decision registered’ event.

The ’hit’ or ’stay’ choices of both players were manually entered by the experimenter. Reaction times could vary based on the experimenter and thus were not further analyzed.

The rule for punishing a player that overshot was introduced to encourage the player who played after an overshooting event to continue playing and not immediately choose ’stay’. This helped to keep the game more dynamic and prevented simpler strategies.

Besides the numbers 1 to 10 in a minority of trials (5%) a ’-2’ or a ’14’ could appear. These additional numbers were also included to prevent simple strategies, like always choosing ’hit’ when the first number of a turn is displayed (Figure 1b).

The structure of the simple jack game allowed us to observe EGS neurons while he himself played the game and while his opponent was doing his or her turn. For all purposes in this analysis we will refer to EGS as player 1 (P1) and his opponent as player 2 (P2).

To simplify the analysis, we split each session in which the game was played into those parts in which player one was active and in which player 2 was active. We then segmented each round into single trials which start with presentation of the current number (cue presentation) and end with the moment at which the decision is entered by the experimenter. The 2000 ms section before each trial begins is termed inter trial interval (ITI).

The time directly after the experimenter entered the player’s decision (post-decision) was confounded since the trial continues differently from that point on. In case of a ’hit’ decision a 2000 ms blank period follows and in case of a ’stay’ decision the score of the current player is displayed. Because of this dichotomy (in one case a visual is displayed and a specific sound for stay is played and in the other nothing is displayed and the sound for hit is played) and the fact that trial progression after the decision was different for different trials, this post-decision period was confounded and not further considered for analysis.

During the task EGS was sitting in his wheelchair 2 m in front of a LCD screen (1190 mm screen diagonal). Stimulus presentation was controlled using a combination of Unity (Unity Technologies, San Francisco, CA) and MATLAB (The Mathworks Inc., Boston, MA). EGS observed the game while the second player, either one of the experimenters or an occupational therapist, performed his/her turn but the observation was not systematically controlled. Sessions where EGS was inattentive or tired were excluded from analyses.

## Analysis

### Epochs

Depending on the events during the course of a trial we selected four distinct epochs for analysis (Figure 1c). The visual epoch was defined by a 400 ms window 100 ms after the onset of the card presentation. From previous experiments and NHP electrophysiology it is known that neurons in the parietal cortex will react to visual stimuli and this epoch would allow us to look into the early neuronal response. Next, we defined a 400 ms window right before the response phase as the delay epoch. During this epoch no visual stimulus was visible on the screen but we assumed that processes associated with decision making would be driving some neurons in this epoch. We termed the third epoch early decision epoch and defined it as a 400 ms window 100 ms after the decision phase onset. Since a red question mark on the screen would visually indicate the decision phase we expect some visual response in this phase in addition to the dynamical response that correlates to the decision-making process. Finally, the late decision epoch was defined as a 400 ms window right before the time when the decision of EGS or the other player was registered. All epochs were chosen to capture the most important moments within the decision-making process based on the events during a single trial.

### Task parameters

To assess different parameters that might be used throughout the simple jack task we defined several parameters and tested each unit’s selectivity based on them.

### Decision selectivity

Since neurons in the posterior parietal cortex have been shown to encode the decision value in various tasks we tested our neuron pool for selectivity for one of the two decisions each player had to make during each trial. We tested each unit’s decision selectivity by performing a t-test with unequal sample sizes for the firing rates in ’hit’ or ’stay’ trials for all epochs and also continuously in 50 ms increments during the entire trial time. A unit was considered selective for the decision if the p-value of the t-test was below 0.05. We performed the test for player 1 and player 2 separately.

### Face value selectivity

Another variable important to the task is the current number value that is shown in the card presentation phase. We assessed a unit’s selectivity for the number in two ways. One is the ’face value selectivity’ which was calculated by using the correlation between the firing rates of a unit and the face values presented in each trial. A unit was considered face value selective if the p-value for the correlation was below 0.05 (no distinction was made between positive and negative correlations).

### High-low value selectivity

Another approach to assess if a unit was selective for the number value presented in the card presentation phase was grouping the values into a high (numbers 6-10; excluding 14) and a low (numbers 1-5; excluding -2) category. The firing rates in the two categories were then assessed using a t-test and units which had a p-value below 0.05 were considered high-low value selective. Note that we did not try to fit tuning curves to assess if units were selective for one specific number. This is due to edge effects (for numbers 1 and 10) and the fact that the amount of trials per individual number value was not high enough to allow for such an approach.

### Cumulative value selectivity

In addition to tracking the current number value we assumed that some units might keep track of the current cumulative score of a player. To evaluate this, we calculated the cumulative value in each trial which is the sum of all face values from the current and all previous trials of a turn. We then calculated the correlation between a unit’s firing rate and the cumulative value. Units with a p-value smaller than 0.05 were considered cumulative value selective. Note that the cumulative value selectivity is somewhat related to the decision value selectivity because a higher cumulative score might be associated with a higher likelihood to choose ’stay’. However, the actual decision is not only based on the cumulative value but also on the score of the opponent (when the other player starts a round) and the risk assessment for the current round (total score). Therefore, both of these task parameters are treated separately.

## Results

### General

Overall, we recorded 500 single units from AIP, which had a minimum mean firing rate of 1.5 Hz during trials, over all 39 games which were recorded. In 13 of the games EGS played against the experimenter and in the remaining 26 games EGS played against one of the occupational therapists. EGS won 49 % (19) of the games. The mean number of trials per game was 55 ± 11 for both players. The mean number of trials per turn was 3.39 ± 0.33 for Player 1 and 3.36 ± 0.31 for Player 2.

### Example units

In the first step of the study we determined multiple task parameters which might be represented by neurons in AIP. Based on these parameters (see Methods section) we traced single units that were specifically selective for them. In Figure 2 we show two separate example neurons from different sessions and channels (both from the AIP array) which exhibit selectivity for two of these parameters. Figure 2a shows the mean spike density over the course of trials in which EGS choose ’hit’ (red line) and ’stay’ (blue line). The firing rates for the two conditions separate already early on in the trial. A distinctive visual response peak can be seen in the card presentation phase. However, the biggest firing rate difference starts to appear after the decision cue (’?’; see Figure 1a) is presented when the firing rate for ’stay’ decisions is about three times higher than for ’hit’.

**Figure 2:**
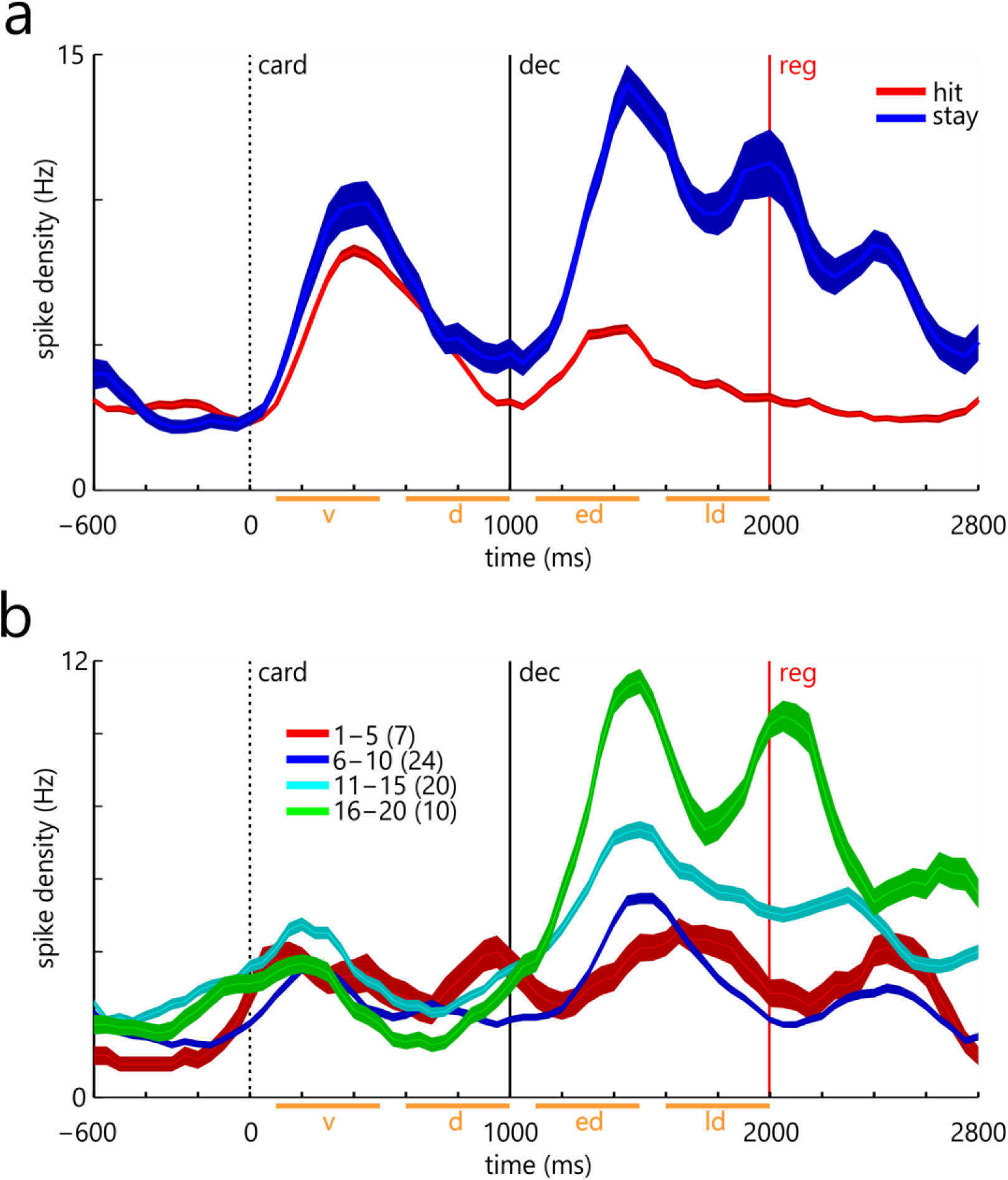
(a) Spike density function of an example neuron which has a high firing rate when EGS decided to choose ’stay’ during the game. Mean spike density for ’hit’ (red) and ’stay’ (blue) trials and their standard error (shaded area). Note that the standard error for stay trials was much higher since ’hit’ was chosen about 3 times more often than ’stay’. Trials were aligned to the card presentation phase (card). The vertical lines represent the three events - card presentation (card), decision (dec) and decision registered (reg) – in a trial. The red vertical line indicates the mean time at which the response of the player was entered by the experimenter which could differ from trial to trial. (b) A different example neuron which shows a graded response for the current cumulative value. The colored lines indicate the mean spike density for all trials for a range of cumulative values (red = 1-5; blue = 6-10; cyan = 11-15; green = 16-20) and the number of trials for each of those categories (in brackets). Same conventions are used as for (a).

Figure 2b shows a different example neuron which illustrates characteristics for cumulative value selectivity. The visual peak in the card presentation phase is much smaller than in the previous example. The mean spike densities for the four different cumulative value groups (1-5 red; 6-10 dark blue; 11-15 light blue; 16-20 green) only separate clearly after the decision cue was presented. Although this example neuron is also selective for the decision (not shown) only a fraction of neurons are selective for both, decision value and cumulative value as will be shown later.

Figure 3 shows examples of the correlation of the firing rates of three neurons and the cumulative card value during the late decision epoch. We can find examples of firing rates that are positively correlated to player 1 (blue) and/or player 2 (red) either only for player one (Figure 3a), only for player two (Figure 3b) or both (Figure 3c). As can be seen, not all neurons that show a significant correlation for one of the player’s cumulative values automatically are also significantly correlated to the other player’s cumulative value. All examples show positive correlations but we also found neurons with negative correlations in all three cases (not shown).

**Figure 3:**
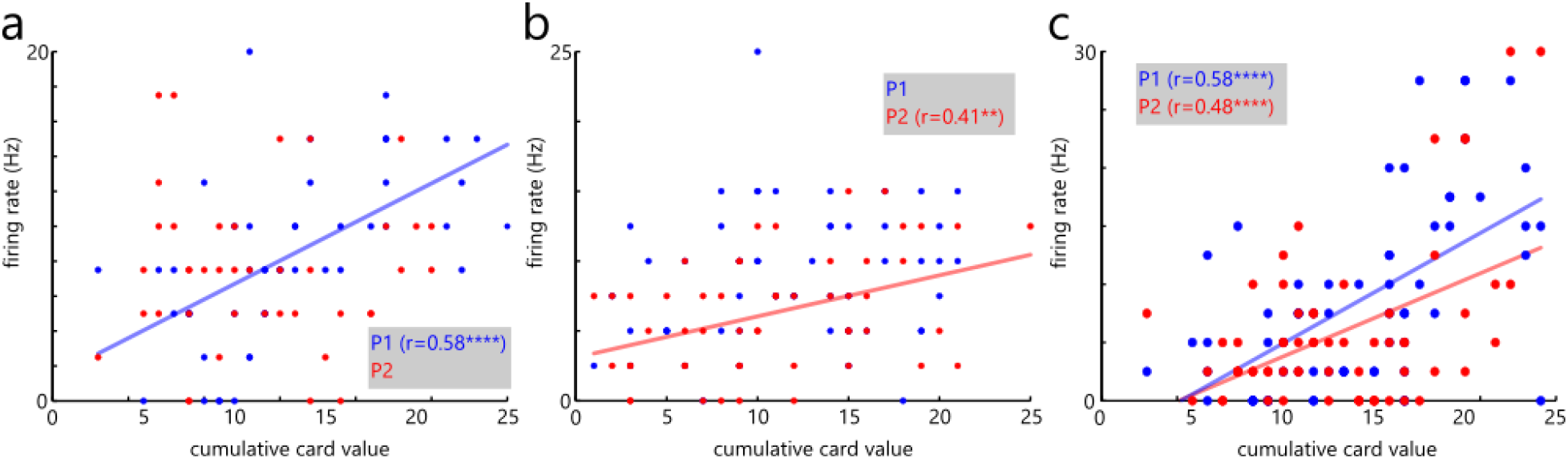
Example correlations of the cumulative value and firing rate for three neurons. One is only correlated to the cumulative value of player 1 (a) the second is only selective for the cumulative value of player 2 (b) and the third one is selective for the cumulative value of both players (c). Blue dots represent cumulative values for player 1 and red dots represent cumulative values for player 2. The red and blue lines are the corresponding regression lines for the two players (only shown for significant correlations). Grey insets show the r-values for significant correlations.

### Population spike density – visual responses

Figure 4a shows the mean spike density for all of the 500 recorded AIP units normalized individually to a time window from -600 to 0 ms before the start of a trial. The normalized unit spike activity was then averaged over the entire population. Two distinct peaks of high activity can be seen in the plot. Their transient nature and proximity to the visual stimuli in the card presentation and decision phase are indicative of a visual response. As has been shown in studies in the posterior parietal cortex of NHPs (Andersen et al., 1987) and our prior human study with the same subject (Klaes et al., 2015) neurons in the posterior parietal cortex are very responsive to visual cues (but not auditory cues as has been shown before in NHPs and humans (Grunewald et al., 1999; Klaes et al., 2015). The average visual response delay calculated from the peak of the responses was 372 ms for the card presentation phase (relative to the card event) and 419 ms for the decision phase (relative to the decision event). The slightly lower peak for the decision phase could be attributed to the somewhat smaller visual stimulus in this phase compared to the card presentation phase.

**Figure 4:**
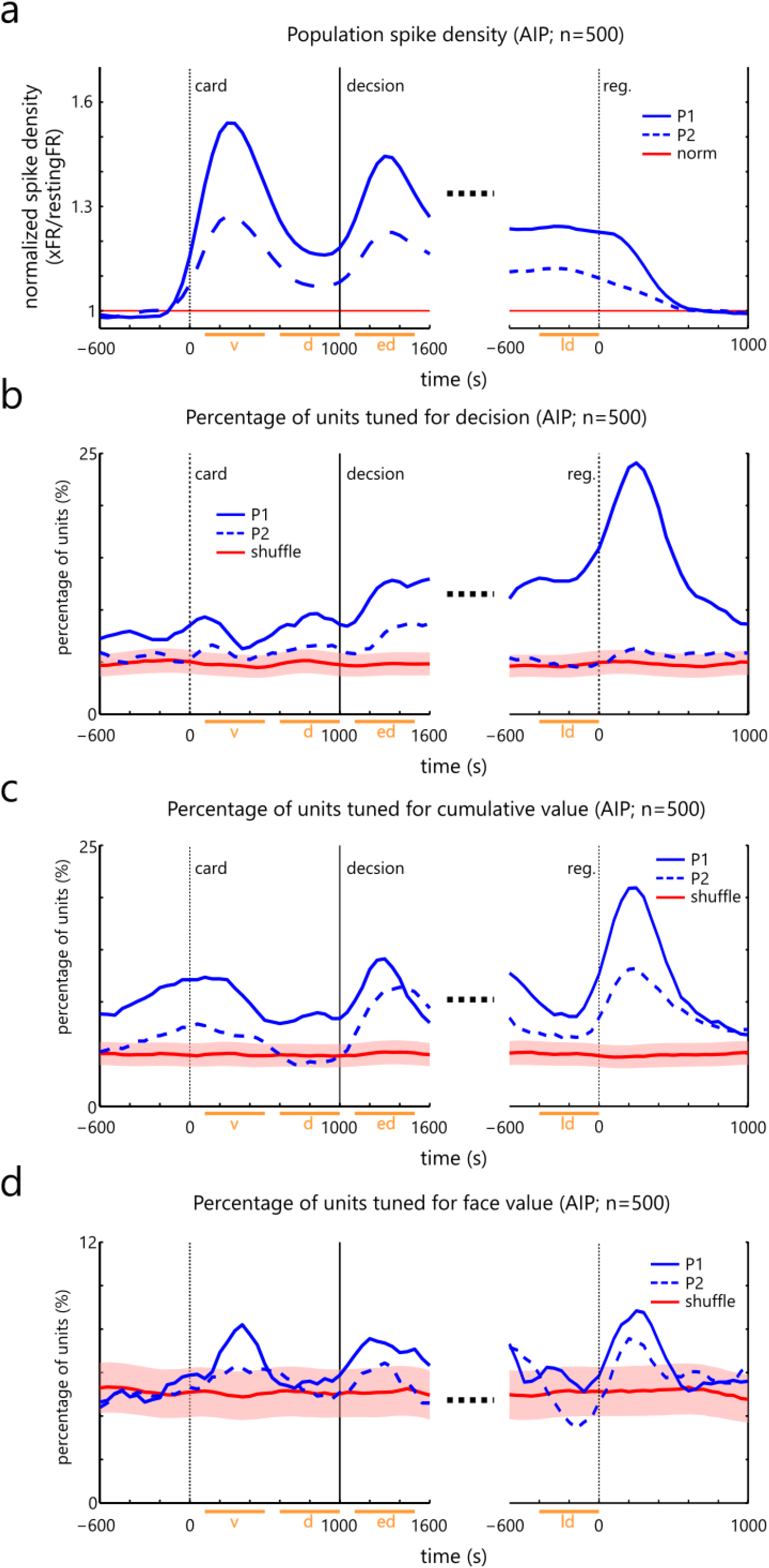
(a) Average spike density and recruitment plots. Mean normalized population spike density. Each unit’s mean spike density is normalized to the first 500 ms right before the card presentation phase (card) and then the mean over all normalized spike densities is taken. Horizontal red line indicates the norm value. Vertical lines show trial events (same conventions as Figure 2). Orange horizontal bars on the time axis indicate epoch times for the visual (v), delay (d), early decision (ed), late decision (ld) epochs. Trials are aligned to the card presentation phase (dotted vertical line at 0 ms). Spike density is separately shown for player 1 (P1) and player 2 (P2). (b) Percentage of units significantly selective for the decision (’hit’ vs. ’stay’) throughout time. Same conventions for time axes are used as in (a). Red line shows result of a shuffle test where decision labels were randomly assigned (shaded area shows standard error). (c) Percentage of units significantly selective for the cumulative value. Same conventions are used as in (b). (d) Percentage of units significantly selective for the face value. Same conventions are used as in (b).

### Recruitment plots

Figure 4b-d shows recruitment plots which show the percentage of neurons from the total recorded population from AIP that showed significant selectivity for one of the task parameters (see Methods). Since there were no significant amounts of neurons with selectivity for the high-low value there are only three recruitment plots shown for the decision selectivity (4b), the cumulative value selectivity (4c) and the face value selectivity (4d). The recruitment plots were calculated by counting the number of neurons in each 50 ms non-overlapping time step throughout the length of a trial beginning 600 ms before card presentation and ending 2800 ms later. Since the registered response of both players could vary from trial to trial only the mean time of this response is shown in the plots (red vertical lines). Also, to take into account units which randomly became significant we also show the mean recruitment for a shuffle test in which the respective task parameter (’hit’ and ’stay’, cumulative value and face value) were randomly assigned (red horizontal lines) together with the standard error of the shuffle tests (shaded red area). Recruitment plots are shown for player 1 (P1; blue solid lines) and player 2 (P2; blue dashed lines) separately.

### Decision selectivity

We found that 39 % (194 units) of all recorded AIP units were significantly selective for the decision of P1 during at least one of the four epochs (see Figure 5a-d; dark grey bars). As expected the number of neurons which are selective for the decision rises during the trial until a maximum is reached around 1800 ms. Interestingly a minority of neurons seems to be selective already before the current card is shown. It turns out however that the number of such neurons is not significantly above what would be predicted from a shuffle distribution (as can be seen in Figure 5a). In later epochs this number becomes highly significant, though (compare Figure 5b-d). This development only holds true for the decision of player 1. The plot shows a rise of neurons that are selective for P2, however, when looking at the number of units selective for the decision in each of the epochs (Figure 5a-d; light grey bars) it becomes clear that these numbers are never significantly above the expectation from the shuffle test. This result is in accordance with what we expected since the decision of player 2 can neither be controlled by EGS nor is it strictly necessary to keep track of it. As we will see this is different for the cumulative value selectivity.

**Figure 5:**
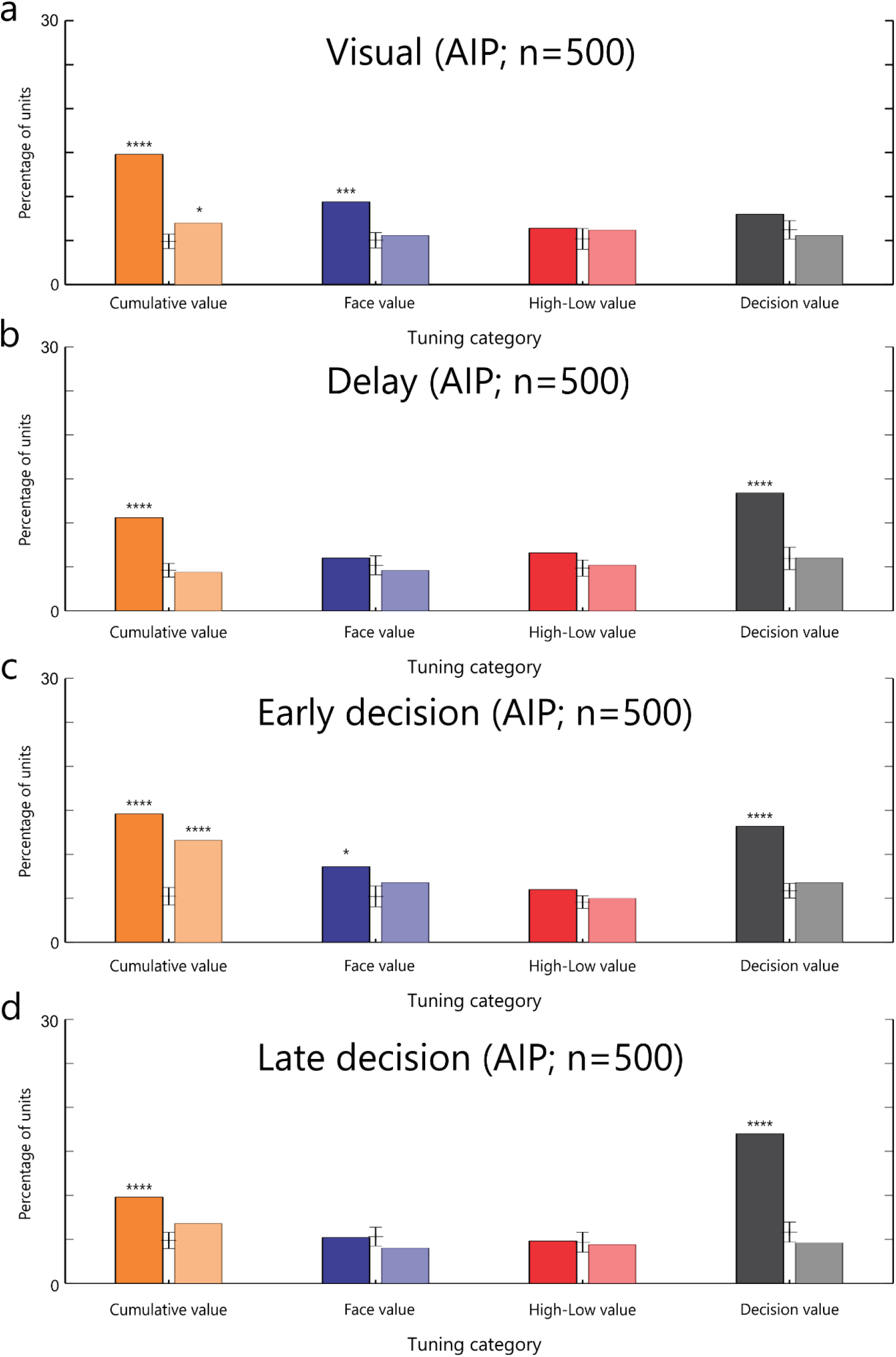
Selective unit statistics. (a) Percentage of units which were selective for specific task parameters during the visual epoch. Each bar shows the number of units that were significantly selective for the four task parameters for player 1 (left bar) and player 2 (lighter right bar). Star notation shows the significance level of the number of units above a shuffle test value in which labels were randomly assigned (small vertical lines between the main bars showing the shuffle mean and standard error). Bar plots for the delay epoch (b), the early decision epoch (c) and the late decision epoch (d) show corresponding values for the different epochs.

### Cumulative value selectivity

Of the 500 recorded AIP neurons 37 % (187 units) were significantly selective for the cumulative value of P1 during at least one of the four epochs. The recruitment plot (Figure 4c) in this case looks very different from the decision recruitment plot. First of all, throughout the whole trial we find a significant number of neurons representing the cumulative value (compare Figure 5; dark orange bars). It seems that at the beginning of the trial we can find a good number of neurons that have a significant selectivity for this task parameter and this number seems to stay up throughout the whole trial. The number of neurons is highest shortly after card presentation and the start of the decision phase (see Figure 5a and 5c compared to 5b and 5d). The highest number is reached in the early decision epoch (see Figure 5c) and goes down after this epoch. This is very different from the decision selectivity recruitment plot where we see a continuous rise of the number of neurons representing the decision. Another difference is the curve for player two (Figure 4c; blue dashed line) which shows a significant number of selective neurons at different times during the trial. Significant numbers of neurons can be found during the visual epoch (Figure 5a; light orange bar) and the early decision epoch (Figure 5c; light orange bar).

### Face value selectivity

Of the total 500 recorded AIP neurons 25 % (126 units) were significantly selective for the face value shown to P1 during two of the four epochs. Unlike the decision selectivity and the cumulative value selectivity we could generally only find a few neurons selective for the face value. Significant numbers above the expectation from a shuffle test could only be seen in the visual epoch (Figure 5a; dark blue bar) and the early decision epoch (Figure 5c; dark blue bar). We also did not see any significant number of neurons selective for face value seen by Player 2 at any time during a trial.

### High-low value selectivity

We did not find any significant numbers of neurons that had a high-low value selectivity in any of the four epochs (Figure 5a-d; dark and light red bars). For this reason, we do not show a recruitment plot for this task parameter.

### Neuron specificity

One important question that is not answered by the recruitment plots is if neurons are selective for only one parameter or multiple ones. To investigate this, we look at the overlap plots in Figure 6. The pie chart for each of the four epochs (Figure 6a-c) show how specific the selective AIP neurons in that epoch are regarding the three primary task parameters (decision selectivity, cumulative value selectivity and face value selectivity; excluding high-low value selectivity since we found no significant numbers of neurons for this parameter). Each pie chart shows the percentage of all neurons that were significantly selective during that epoch and what they were selective for. The solid colors show the percentage of neurons that were only selective for one of the three task parameters, decision selectivity (D; grey), cumulative value selectivity (C; orange) and face value selectivity (F; blue). The striped areas represent neurons which are tuned for two of the task parameters, cumulative/decision (CD; orange/grey), face/decision (FD; blue/grey) and face/cumulative (FC; blue/orange). The black sector shows the percentage of neurons which are selective for all three parameters. It is interesting to note that large proportions of selective neurons are very specific for only one task parameter. Neurons which are selective for multiple task parameters only make up between about 1/4 and 1/3 of the total number of selective neurons. The biggest overlap seems to be decision and cumulative value selectivity which are also the two task parameters which are in general represented most by the entire population and are more similar than the face value selectivity.

**Figure 6:**
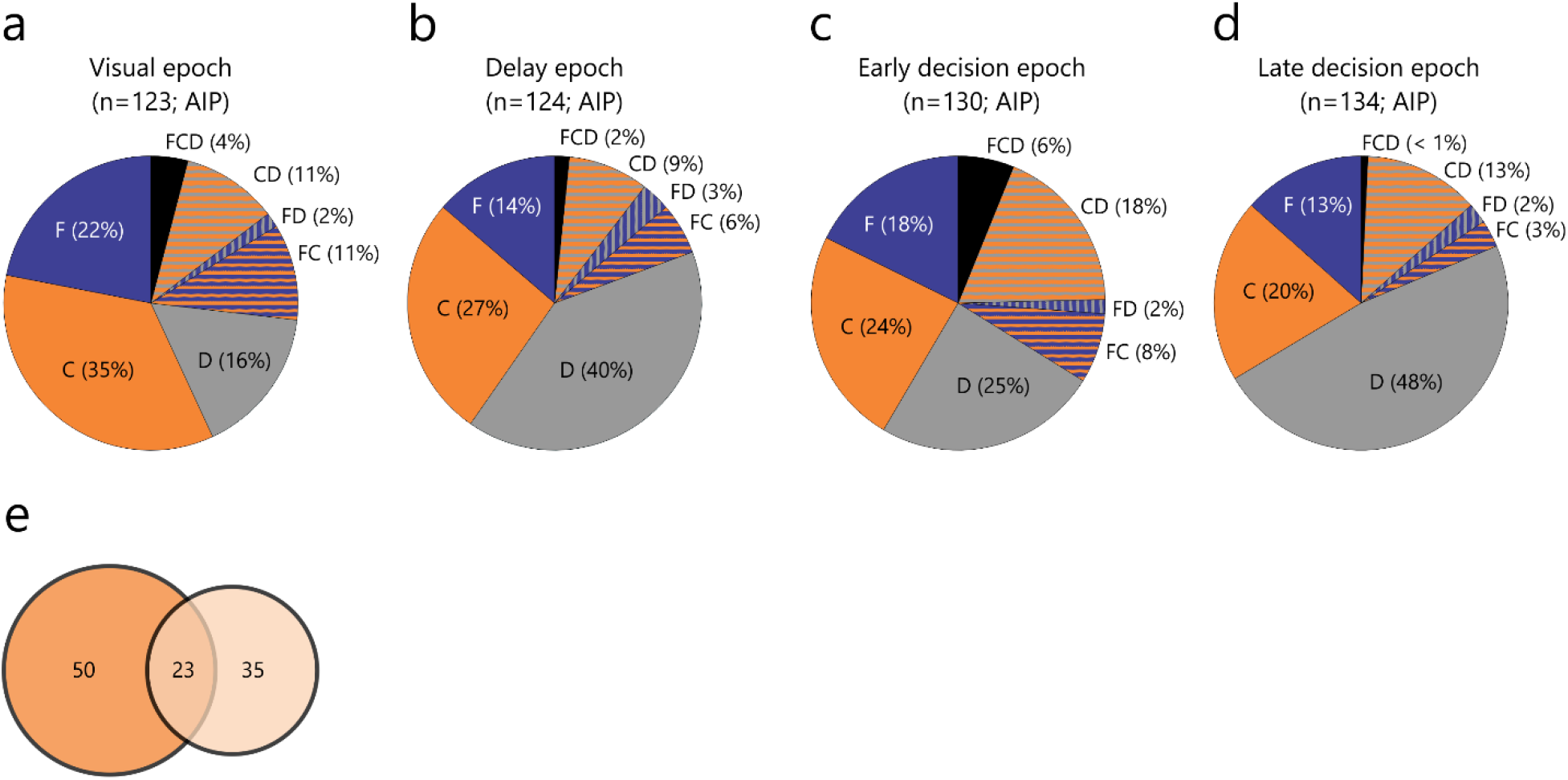
Overview of overlap between units which were significant for the three task parameters during the four distinctive epochs (a-d) as percentage of the total number of units tuned during that epoch. Solid colors show units that were exclusively tuned for the face value (F; blue), the cumulative value (C; orange) and the decision (D; grey). Striped sections show units colored for two of the parameters, face and cumulative (FC; blue/orange), face and decision (FD; blue/grey) and cumulative and decision (CD; orange/grey). Black (FCD) indicates units that were significant for all three parameters. For the early decision epoch the overlap between units selective for the cumulative value for player 1 (solid orange) and player 2 (light orange) is shown as a Venn diagram (e). Note that the numbers in the circles are absolute numbers and not percentages in (e).

A comparison of the overlap between the selectivity for a parameter in player 1 versus player 2 only made sense for the cumulative selectivity. None of the other parameters showed a significant number of selective neurons for player 2 (Figure 5; light colored bars). Since the number of neurons tuned for the cumulative value of player 2 was largest in the early decision epoch, we only show the overlap for this epoch as a Venn diagram in Figure 6e. The overlap shows that a majority of neurons are either exclusively tracking the cumulative value for player 1 or for player 2. Less than a quarter of neurons (21%) are selective for both players.

## Discussion

Studying the neuronal processing of numeric variables in the posterior parietal cortex of a tetraplegic patient showed several interesting results.

First, to our knowledge for the first time we report that single neurons in the human posterior parietal cortex can represent a high-level economic decision, the current cumulative value and the current face value in a complex economic task (two-player, press-your-luck game). It seems that the neurons we observed are involved in multiple functions of the decision process while most of them are most selective for only one of those we tested. Neurons are dynamically recruited at different times during the trial depending on the decision variable in question.

In our task, we did not control for eye movements and general movements of the head or face so it is possible that some of the selectivity of neurons is actually related to those movements during parts of the early decision epoch and especially late decision epoch. However, the cues presented to the patient were all stationary and located in the center of the screen so we would not expect any directional components of these movements. The second spike density peak in Figure 4a would also be more consistent with the idea of visual responses than some effect of muscle activation since it has the same response time relative to the cue event (dec) as the clearly visual response to the card presentation (card). We therefore do not think that our results have been influenced much by these movements.

### Decision selective neurons

The recruitment plot for the decision selective neurons (Figure 4b) shows a steady rise of neurons that are selective for the decision especially after the onset of the decision phase. The maximum number of selective neurons is reached at about 1800 ms after presentation of the current number card or in other words just before the decision was registered (late decision epoch; see Figure 5d). This is compatible with many observations in NHP studies in which a build-up of firing rates just before the action execution can be seen (Kiani and Shadlen, 2009). In our case, increased firing rates relative to baseline would result in increased recruitment of neurons. This view is also compatible with evidence accumulator models, which are the basis of many theoretical mechanisms for perceptual decision making (Ratcliff et al., 1999; Mazurek et al., 2003; Kiani and Shadlen, 2009; Westendorff et al., 2010). However, as has been discussed by Shadlen and Shohamy (Shadlen and Shohamy, 2016) the same framework that is involved in perceptual decision making might also be involved in decisions which are based on memory retrieval. In our case the retrieval would be remembering the current number and the current cumulative value.

### Cumulative value selective neurons

To inform the current decision, the cumulative value has to be stored and later retrieved from (episodic) working memory. The involvement of the lateral posterior parietal cortex in this working memory maintenance (Constantinidis and Steinmetz, 1996) overlaps with its role in other decision-related processes in the human brain such as reward expectancy (Platt and Glimcher, 1999) and action planning (Andersen et al., 1987). Other brain structures, for example the hippocampus and the medial temporal lobe, were also shown to be involved in this putative decision-making circuitry (Davachi, 2006). Although the cumulative value and the decision are related, they represent different parts of the decision process which are apparently utilized at different times as can be seen in the recruitment plots (Figure 4 b,c). It is also interesting to note at this point that the neuronal populations which are selective for the cumulative value (and also for the other variables we investigated) are not completely overlapping between the two players. In addition to that these multifunctional neuron populations had different functions in other tasks which were described by our previous studies (Aflalo et al., 2015; Klaes et al., 2015). This behavior has also been termed ‘mixed selectivity’ and can explain many of the observed neuronal characteristics (Fusi et al., 2016). This mixed selectivity can be seen for the various decision variables we investigated in this study and the overlapping neuronal populations which are selective for them. In addition, it can also be seen in regard to completely different tasks when compared to the previous studies.

### Face value selective neurons

Interestingly we did not find many neurons which were selective for face value. Those few that were selective for face value were only selective during specific times during the trial. One reason for the low number of face value selective neurons could be our definition of face value selectivity. If the neurons do not encode the face value linearly as is assumed by our selection definition (see Methods), for example because they have a narrow selectivity for a specific number, they would not be detected by our definition. Since the Simple Jack game did not provide for enough trials to generate statistically sound tuning curves for single numbers we cannot exclude this possibility. To address this issue an explicit study would be needed which tests various number values which were outside the scope of the Simple Jack game. However, we could find significant numbers of neurons tuned for face value by our definition in the visual and the early decision periods. The reason for this could be that these are the two periods when the current number is highly relevant. In the visual epoch the current number would be especially important because that is when the current number is presumably identified. In the early decision epoch the current number would be important because this is presumably when the numbers are added up and compared to the cumulative value. This finding suggests that mental processing of numbers for an upcoming economic decision follows a timeline in which a number’s face value is transformed into a more complex value informing the decision. Such mental operations have been previously demonstrated to critically involve human intraparietal areas (Knops et al., 2009).

### Reduced neuronal activity for P2 vs P1

The dynamics of tuned units for P2 seem to generally follow the dynamics for P1 for all three task parameters which we investigated. When significant numbers relative to the shuffle test are considered for the four task epochs we only found significant numbers of specific neurons for the cumulative value and only in some of the epochs. This is in line with prior finding, showing intraparietal involvement in observing actions of others both in NHP (Fogassi et al., 2005; Rozzi et al., 2008; Bonini et al., 2010) and humans (e.g. Chong et al., 2008).

One possibility why the numbers for P2 are lower could be that EGS did not always pay attention (e.g. to the monitor) when his opponent was playing. AIP activity was reported to be modulated by attention (e.g. Peck et al., 2009) and, likewise, not attending the game would lead to overall lower numbers of selective units. Since we also know that the neuron populations which are selective for task parameters of P2 are just partially overlapping with P1 this view could be wrong. The population of neurons selective for P2 task parameters could also be lower because the opponent’s behavior in the game is arguably less important for success than one’s own. This would be especially true for the decision of the opponent, which cannot be influenced by the other player at all. For the same reason the current face value that the opponent is seeing would be less important for the player. On the contrary, since the cumulative value the opponent achieved during his or her round has to be tracked to make a good decision it would be useful to have this information. As it turns out the cumulative value is the only decision variable we investigated where significant numbers of neurons can be found in some epochs of the task for P2. This supports the hypothesis that neurons in the parietal cortex are only selective for those decision variables which are relevant for the task at hand. This means that AIP neurons might represent the outcome of number operations informing the current decision-making process (and relevant to the current situational context) rather than the numbers alone. While our data suggests this might be the case, this hypothesis needs further testing.

In summary, the three task parameters: face value, cumulative value and decision show very specific dynamics during the trial. The highest numbers of selective AIP neurons can be found for the decision and the cumulative value. On the one hand, it seems that relevant information is available at critical points during a trial and partially overlapping populations of neurons are dynamically selective for these variables. This includes tracking specific task variables during the other player’s turn. On the other hand, task parameters that are not task relevant during the opponent’s turn (such as the face value) seem to be only weakly if at all represented by AIP.

In comparison to our previous studies with the same patient and with the same electrode configuration (Aflalo et al., 2015; Klaes et al., 2015) we find that neuron populations in the same areas of AIP have been variably used to represent task relevant parameters for different tasks. For example, neurons recorded from exactly the same electrodes were used to determine various hand shapes in a rock-paper-scissors type of task (Klaes et al., 2015) or in various brain computer interface scenarios in which hand and arm operations were imagined (Aflalo et al., 2015). This again is compatible with the idea that high level cortical areas contain non-linear mixed selectivity neurons which are not tuned for one particular feature but which are representing variable task parameters depending on a given task (Fusi et al., 2016). Especially in comparison to other tasks in which EGS had to imagine arm or hand movements it is important to stress again that the Simple Jack task did not require any such motor imagination. Still, we found neurons which correlated with EGS’ economic decision in the Simple Jack game although this decision did not relate to any instructed imagined movement. This is, to the best of our knowledge, the first demonstration of human economic decisions being possible to track on a single neuron level. Future studies will be needed to investigate in more detail how the specific dynamics of selectivity come about and how single numbers and their transformations are represented by narrowly selective neurons. This study can be seen as a groundwork to determine the scope of more specific experiments of complex numerical operations and economic decisions on a single neuron level.

## Acknowledgements

This work was funded by the Deutsche Forschungsgemeinschaft (DFG, German Research Foundation), project ID 122679504, SFB 874/A11, NIH (grants EY013337, EY015545 and P50 MH942581A), the Boswell Foundation, the T & C Chen Brain-machine Interface Center at Caltech, and the USC Neurorestoration Center. We thank Viktor Shcherbatyuk for computer assistance, Tessa Yao, Alicia Berumen, and Sandra Oviedo for administrative support, and Kirsten Durkin for nursing assistance.

